# Synaptic proteins expressed by oligodendrocytes mediate CNS myelination

**DOI:** 10.1101/410969

**Authors:** Alexandria N. Hughes, Bruce Appel

## Abstract

Oligodendrocytes ensheath neuronal axons with myelin, a proteolipid-rich membrane that increases conduction velocity and provides trophic support. Our lab and others have provided evidence that vesicular release from neurons promotes myelin sheath growth. Complementarily, transcriptomic and proteomic approaches have revealed that oligodendrocytes express many proteins that allow dendrites to sense and respond to vesicular release at synapses. Do axon-myelin contacts use similar communication mechanisms as nascent synapses to form myelin sheaths on axons? To test this, we used fusion proteins to track synaptic vesicle localization and membrane fusion within spinal cord axons of zebrafish larvae during developmental myelination. Additionally, we used a CRISPR/Cas9-mediated GAL4 enhancer trap and genetically-encoded intrabody to detect expression and localization of PSD-95, a component of dendritic postsynaptic complexes, within oligodendrocytes. We found that synaptic vesicles accumulate at ensheathment sites over time and are exocytosed with variable patterning underneath myelin sheaths. Accordingly, we also found that most, but not all sheaths localized PSD-95 with patterning similar to exocytosis site location within the axon. By querying published transcriptome databases, we found that oligodendrocytes express numerous transsynaptic adhesion molecules that function across synapses to promote synapse formation and maturation. Disruption of candidate PDZ-binding transsynaptic adhesion proteins in oligodendrocytes revealed that these proteins have variable effects on sheath length and number. We focused on one candidate, Cadm1b (SynCAM1), and demonstrated that it localized to myelin sheaths where both its PDZ binding and extracellular adhesion to axons are required for myelin sheath growth. Our work reveals shared mechanisms of synaptic and myelin plasticity and provides new targets for mechanistic unraveling of activity-regulated myelination.

## INTRODUCTION

Communication in the central nervous system (CNS) depends on billions of connections. Synapses, the connections between neurons, are plastic structures that grow and change with experience-evoked neuronal activity. Additionally, connections form between neurons and oligodendrocytes, the myelinating cell type of the CNS. Myelin sheaths also are plastic structures, mutable in length, number, and thickness^1–3^. Some of this plasticity may be triggered by experience, because motor learning paradigms increase myelination^4,5^ and social and sensory deprivation paradigms reduce myelination of relevant brain regions^2,6,7^. What accounts for myelin sheath plasticity? One possibility is that neuronal activity tunes ensheathment, similar to activity-dependent plasticity at synapses^8^. Consistent with this possibility, Demerens and colleagues first demonstrated more than 20 years ago that inhibiting action potential propagation with tetrodotoxin (TTX) reduced myelination of cultured neurons^9^. Activity-mediated communication from axons to myelin sheaths could ensure that active neurons receive enough myelin to faithfully propagate impulses toward termini, facilitating circuit function and higher order cognitive processes^10^. However, almost nothing is known about communication between axons and myelin sheaths.

Neuronal vesicular release promotes sheath growth and stability^11–13^. Upon cleavage of neuronal vesicular release machinery by clostridial neurotoxins, oligodendrocytes formed shorter sheaths on impaired axons and selected unimpaired axons when presented with a choice^13^. Importantly, timelapse imaging revealed that oligodendrocytes retract nascent sheaths from impaired axons more quickly than from controls^11^. These studies indicate that oligodendrocyte processes can detect and respond to axonal vesicular secretion. How might oligodendrocytes detect vesicular release? Oligodendrocytes express numerous genes encoding neurotransmitter receptors, synaptic scaffolds, transsynaptic adhesion molecules, and Rho-GTPases^14–16^ that, when expressed by neurons, endow dendrites with the ability to detect and respond to release at synapses. Similar to their function in dendrites, these proteins may allow oligodendrocytes to sense, adhere, and coordinate a morphological response to axonal secretion^17^. Do oligodendrocytes utilize these proteins similarly to neurons to stabilize nascent myelin sheaths on axons?

Determining how neurons communicate to oligodendrocytes to shape developmental myelination requires an experimental model where both axons and oligodendrocytes can be monitored and manipulated during normal myelination. Here, we examined features of synapse formation between axons and their myelinating oligodendrocytes, including the accumulation of presynaptic machinery, exocytosis at the junction, and postsynaptic assembly. These synaptic features manifest at the axon-glial junction from the onset of zebrafish myelination at 3 days post fertilization (dpf) through 5 dpf, when sheaths are stabilized. Surprisingly, we uncovered unforeseen diversity in the synaptic features present at these contacts: exocytosis sites varied in shape and position under sheaths, and the major postsynaptic scaffold PSD-95 also localized with variable position within some, but not all sheaths. To test whether the synaptic features of the axon-myelin interface are important for normal myelination, we manipulated synaptogenic adhesion proteins in oligodendrocytes. When these same proteins are disrupted in neurons, synaptogenesis falters and synapses are abnormal in size and number. Analogously, oligodendrocytes expressing dominant-negative adhesion proteins formed myelin sheaths with abnormal length and number. We focused on one candidate, Cadm1b (SynCAM1), and found that it localized to myelin sheaths, where its PDZ binding motif is required for myelin sheath growth. Furthermore, by manipulating the extracellular domain of Cadm1b we found that transsynaptic signaling to axons promotes myelin sheath length. Our work reveals shared mechanisms of synaptic and myelin plasticity and raises the possibility that synaptogenic factors have previously unrecognized roles in developmental myelination.

## RESULTS

### Axons accumulate functional synaptic vesicle machinery under nascent myelin sheaths

Our lab and others have produced evidence that vesicular secretion from axons promotes myelin sheath growth and stability. However, we know nothing about the temporal or spatial qualities of vesicular communication to oligodendrocytes. To determine where vesicular release occurs to support sheath growth, we first investigated the localization of vesicular release machinery in axons that are myelinated in an activity-regulated manner^11^. By imaging live *Tg(sox10:mRFP; phox2b:GAL4; UAS:syp-eGFP)* and *Tg(sox10:mRFP; phox2b:GAL4)* larvae transiently expressing *UAS:*Vamp2-eGFP we tracked the synaptic vesicle proteins Synaptophysin (Syp) and Synaptobrevin (Vamp2) in individual *phox2b*+ axons (Fig. 1A,B). At early larval stages, *phox2b*+ axons are sparsely covered by myelin sheaths (∼15% of length)^11^, which permitted us to visualize synaptic vesicle machinery in both ensheathed and bare stretches of these axons. We found Syp-eGFP distributed in axons in a clearly punctate pattern (Fig. 1B), whereas Vamp2-eGFP diffusely labeled membrane and obscured intracellular puncta (Fig. 1B’), similar to previous observations^18^. We quantified Syp-eGFP puncta, defined as line profile fluorescence peaks exceeding 150% of inter-puncta fluorescence, over the course of developmental myelination in both bare and ensheathed regions. At every stage of development, the number of these puncta correlated positively with sheath length (Fig. 1C), consistent with previous work indicating a positive influence of synaptic vesicle release on sheath growth^11–13^. By 4 days post fertilization (dpf), and continuing through 5 dpf, axons accumulated more Syp-eGFP puncta at ensheathment sites relative to bare regions of axon (Fig. 1D). These data indicate that axons cluster synaptic release machinery at ensheathment sites over time, consistent with the possibility that vesicular secretion mediates axon-oligodendrocyte communication.

**Figure 1.**
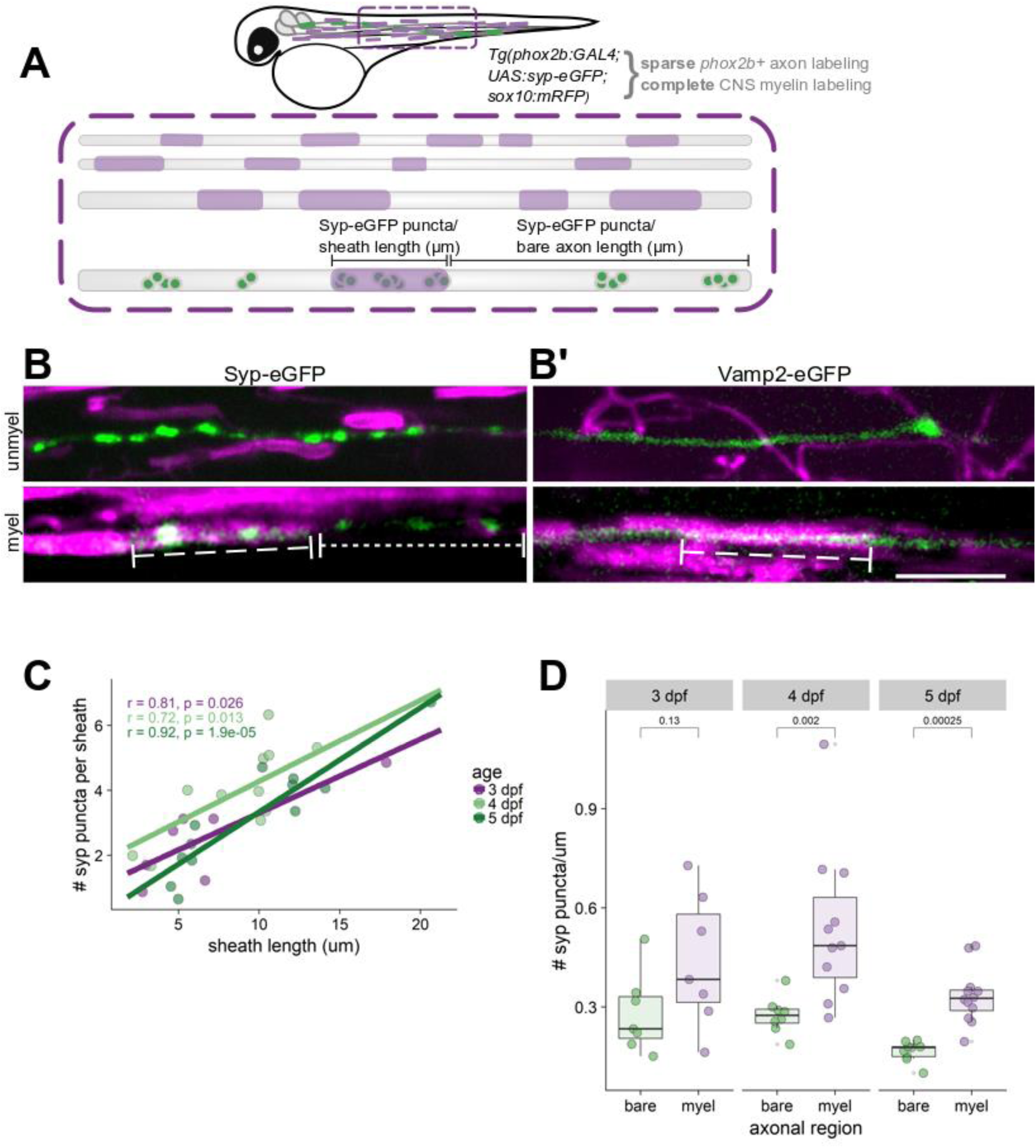
Axons accumulate synaptic vesicle release machinery under myelin sheaths. (A) Schematic of transgenes used to assess synaptic puncta density along myelinated and bare regions of axons. (B) Image of Syp-eGFP in unmyelinated (top) and myelinated (bottom) *phox2b*+ axons in a living 4 dpf *Tg(phox2b:GAL4; UAS:syp-eGFP; sox10:mRFP)* larva. Bracketed dashed lines indicate myelin sheaths; dot-dashed lines indicate bare regions of axons. Scale bar, 10 µm. (B’) Same as (B) for Vamp2-eGFP; animal genotype is *Tg(phox2b:GAL4; sox10:mRFP)* with transient expression of *UAS*:Vamp2-eGFP. Scale bar, 10 µm. (C) Number of axonal Syp-eGFP puncta plotted vs sheath length for sheaths at 3, 4, and 5 dpf. n=7 sheaths/18 puncta (3 dpf); 11 sheaths/43 puncta (4 dpf); 12 sheaths/38 puncta (5 dpf). Pearson’s correlation values (r) are provided for each age. (D) Comparisons of Syp-eGFP density within bare vs myelinated regions of axons at each timepoint. n=7 bare regions/98 puncta and 7 sheaths/18 puncta (3 dpf); 8 bare/119 puncta and 11 sheaths/43 puncta (4 dpf); 9 bare/75 puncta and 12 sheaths/38 puncta (5 dpf). Wilcox rank-sum test, p=0.13, p=0.002, p=0.00025 for 3, 4, and 5 dpf, respectively.

If axons cluster synaptic vesicle machinery at ensheathment sites, is that machinery functional? To test this, we expressed an exocytosis reporter, Syp-pHluorin (SypHy)^19^, in reticulospinal axons. Because pHluorin fluorescence is pH-sensitive, SypHy fluorescence is quenched within acidic synaptic vesicles but fluoresces upon exposure to the neutral pH of the extracellular space. Following endocytosis and vesicle reacidification, SypHy signal is again quenched (Fig. 2A). By pharmacologically inhibiting vesicle reacidification with the v-ATPase inhibitor bafilomycin A1, SypHy signal becomes detectable after an initial exocytic event but is not quenched upon later endocytosis, allowing identification of exocytosis “hotspots” along axons^20^ (Fig. 2B). We used this strategy to ask whether exocytosis hotspots exist under myelin sheaths. We raised embryos microinjected with *Tol2.neuroD:sypHy* DNA and mRNA encoding Tol2 transposase at the 1-cell stage until 4 dpf, the timepoint at which we first detected enhanced Syp-eGFP clustering under sheaths (Fig. 1D). At 4 dpf, we treated larvae with 1 µM bafilomycin or DMSO vehicle and allowed them to behave freely for 1 hour before anesthetizing and mounting for imaging. SypHy fluorescence was minimal when larvae were exposed to DMSO vehicle, reflecting only the net balance of exo- and endocytosis along axons (Fig. 2A’). By contrast, bafilomycin treatment revealed numerous exocytic hotspots along reticulospinal neurons, reflecting net exocytic activity over the hour (Fig. 2B’).

**Figure 2.**
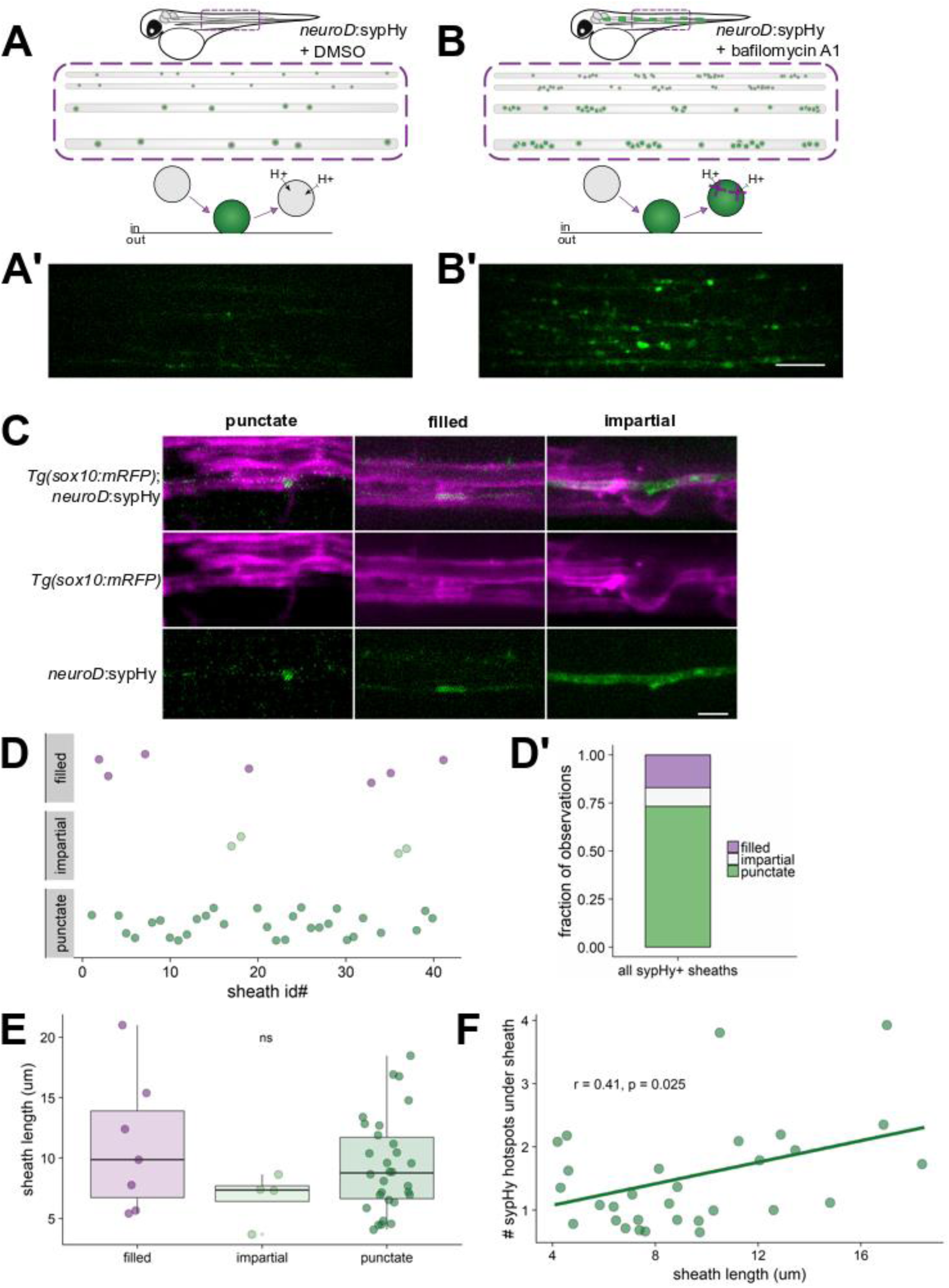
Variable synaptic vesicle exocytosis sites under myelin sheaths. (A) Schematic of Syp-pHluorin (SypHy) function and transient expression in spinal cord neurons, followed (A’) by a live-image of a 4 dpf larva expressing *neuroD*:sypHy. Scale bar, 10 µm. (B, B’) Same as (A, A’) but in the presence of bafilomycin A1 to inhibit vesicle reacidification. Note appearance of several SypHy+ “hotspots” along axons. Scale bar, 10 µm. (C) Expression of *neuroD*:sypHy in *Tg(sox10:mRFP)* larvae treated with bafilomycin reveals three types of SypHy-reported exocytosis sites under myelin sheaths: punctate, filled, and impartial. Scale bar, 5 µm. (D) Raw classification of 41 sheaths by category: 30 punctate, 7 filled, 4 impartial. Presented as a bar in (D’) for visible contribution of each category to the total. (E) No association between category and sheath length, Kruskal-Wallis test. (F) Among the punctate category, the number of SypHy hotspots (detected by same method as Syp-eGFP) was positively associated with sheath length, Pearson’s correlation.

We used our SypHy/bafilomycin paradigm to look for hotspots of exocytosis under myelin sheaths. To do so, we performed our microinjections into *Tg(sox10:mRFP)* embryos at the 1-cell stage and imaged bafilomycin-treated larvae at 4 dpf. SypHy hotspots were abundant underneath myelin sheaths (Fig. 2C) but were variable in morphology. Frequently, SypHy hotspots were punctate, resembling the size and shape of the Syp-eGFP fusion protein (Fig. 1B) and puncta were almost exclusively located near one or both ends of the sheath. In addition, some hotspots were restricted to the ensheathed region but diffusely bright underneath the sheath (“filled”), and occasionally we found axons that were illuminated along their entire length with no visually discernable differences under myelin sheaths (“impartial”) (Fig. 2C,D). No category (punctate, filled, impartial) was associated with longer sheaths than other categories (Fig. 2E), suggesting that the diversity of release site shape or position does not account for differences in sheath length. However, among sheaths with punctate hotspots, the number of hotspots per sheath was positively associated with sheath length (Fig. 2F), corroborating the correlation between accrued synaptic release machinery and sheath length that we found with Syp-eGFP (Fig. 1C). Furthermore, these data complement previous work from our lab and others demonstrating that neurotoxin-mediated cleavage of release machinery reduces sheath growth^11–13^ and are consistent with endogenous vesicle fusion promoting sheath length.

### Myelinating oligodendrocytes express the major postsynaptic scaffold PSD-95 and localize it to sheaths

If ensheathed axonal regions acquire characteristics of presynaptic terminals, do oligodendrocyte sheaths share features of postsynaptic terminals? We investigated the oligodendrocyte expression and localization of the membrane-associated guanylate kinase (MAGUK) postsynaptic density protein 95 (PSD-95), a scaffold known for anchoring neurotransmitter receptors at excitatory postsynaptic terminals. To detect PSD-95 expression in cells of the spinal cord, we used a CRISPR/Cas9-mediated GAL4 enhancer trap strategy^21^ to generate a knock-in transgenic animal that reports expression of *dlg4b*, the gene encoding PSD-95. When we crossed *Tg(dlg4b:GAL4; UAS:eGFP-CAAX)* and *Tg(mbpa:tagRFP)* adults, we generated larvae expressing eGFP-CAAX in *dlg4b*+ cells and tagRFP in *mbp*+ myelinating oligodendrocytes (Fig. 3A). Most *mbp*+ oligodendrocytes also expressed *dlg4b*, indicating that myelinating oligodendrocytes of the spinal cord express PSD-95. As expected, we found that many *dlg4b*+ cells do not express *mbp*, likely reflecting expression of PSD-95 in neurons.

**Figure 3.**
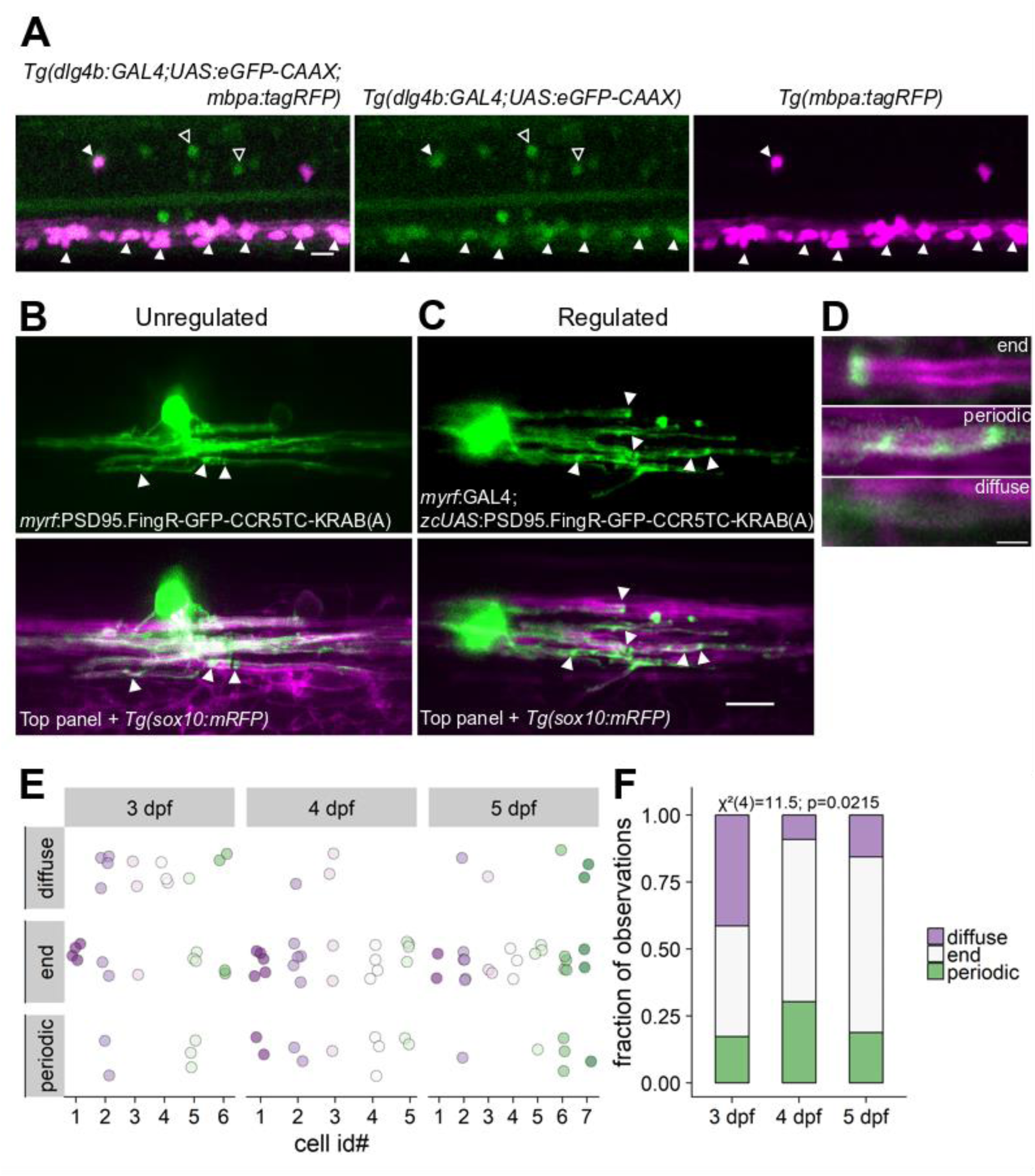
PSD-95 is expressed by myelinating oligodendrocytes and is variably localized within myelin sheaths. (A) CRISPR/Cas9-mediated GAL4 enhancer trap of *dlg4b* reports spinal cord cells that express *dlg4b* via *UAS*:eGFP-CAAX expression. Larvae additionally carrying the *mbpa*:tagRFP transgene reveal *mbp*+ myelinating oligodendrocytes. Closed arrowheads indicate *mbp*+, *dlg4b*+ oligodendrocytes, whereas open arrowheads indicate *mbp*-, *dlg4b*+ cells (likely neurons). Scale bar, 10 µm. (B) Expression of PSD95.FingR to detect PSD-95 localization in individual oligodendrocytes under *myrf* regulatory DNA without a zf-binding site for transcriptional repression. (C) Implementation of transcriptional regulation by providing a zf-binding site upstream of *UAS* (*zcUAS*), driven by *myrf*:GAL4. For both (B) and (C), arrowheads show puncta, scale bar 10 µm. (D) Expression of the transcriptionally-regulated system with *Tg(sox10:mRFP)* to label myelin shows that not all sheaths display localized PSD-95 puncta (“diffuse”), and among those that do, labeling is either restricted to the sheath end (“end”), or periodic along the length of the sheath (“periodic”). Scale bar, 2 µm. (E) Raw classification plot of sheath labeling patterns (y-axis) for individual cells (x-axis) at 3, 4, and 5 dpf. n=6 cells/29 sheaths (3 dpf); 5 cells/33 sheaths (4 dpf); 7 cells/32 sheaths (5 dpf). Note that most cells had sheaths with different labeling patterns. Over developmental time, the distribution of labeling patterns changes by Chi square test (F), notably with a reduction of diffuse labeled sheaths, but all three categories persist at 5 dpf.

To detect the localization of endogenous PSD-95 in oligodendrocytes, we expressed a genetically-encoded intrabody developed by Gross et al (2013) that binds zebrafish PSD-95^22,23^. The intrabody, termed PSD-95.FingR, is fused to GFP and contains a transcriptional regulation system to allow unbound FingR to repress further transcription. This regulatory system requires a zinc finger (zf) binding site upstream of the regulatory DNA driving expression of the construct. We first expressed the FingR in oligodendrocytes directly with putative regulatory DNA upstream of the *myrf* gene and without a zf binding site (“unregulated”). We found that unregulated PSD-95.FingR-GFP broadly labeled the cytoplasm of oligodendrocytes, with some puncta evident at the ends of sheaths (Fig. 3B, arrowheads). We then implemented the transcriptionally-regulated system by providing a zf binding site upstream of *UAS* sequence^23^ that drives PSD-95.FingR expression via co-expression of *myrf*:GAL4 (Fig. 3C). In contrast to the unregulated system, transcriptional regulation of the intrabody unveiled several PSD-95 puncta within most, but not all sheaths (Fig. 3C, arrowheads).

To determine the subcellular/subsheath localization of PSD-95, we next assessed the localization of intrabody signal within sheaths using the regulated form of PSD-95.FingR in a *Tg(sox10:mRFP)* line at 3, 4, and 5 dpf. We identified three unique patterns of sheath labeling: puncta restricted to ends of sheaths (“end”), periodic puncta along the length of sheaths (“periodic”), and faint, diffuse labeling with no obvious localization (“diffuse”) (Fig. 3D). Most oligodendrocytes had sheaths with variable labeling: some sheaths contained “periodic” puncta, while other sheaths of the same cell exhibited only “end” puncta. Our raw observations of puncta locations are plotted in Fig. 3E, with individual oligodendrocytes on the x-axis and all discernable sheath puncta types plotted per oligodendrocyte on the y-axis. The most frequent labeling pattern was “end” puncta at every age we examined. However, the distribution of these localization patterns changed over the developmental period of myelination (Fig. 3F), notably via a reduction in diffusely-labeled sheaths. Taken together, these data suggest that some, but perhaps not all, oligodendrocyte sheaths accumulate PSD-95 over the course of development.

### Synaptogenic adhesion molecules variably tune sheath length and number

In neurons, PSD-95 anchors receptors, ion channels, and synaptic signaling molecules at the postsynaptic membrane via its PDZ domains. These domains are found in PSD-95 and other synaptic scaffolding proteins, where they bind short, C-terminal PDZ binding motifs located on target proteins destined for synapse localization. PDZ binding motifs direct the localization and function of many synaptogenic, transsynaptic adhesion proteins. This family includes members of the neuroligins, synCAMs, netrin-G ligands, and the leucine rich repeat transmembrane proteins^24^. In addition to adhering pre- and postsynaptic terminals, these proteins are potent signaling molecules. The signaling they induce across the nascent cleft is sufficient to induce synaptogenesis even when ectopically expressed in non-neuronal HEK293 cells: cocultured neurons assemble presynaptic terminals and release neurotransmitter onto expressing HEK293 cells^25–28^. Intriguingly, many of these synaptogenic adhesion molecules are expressed by oligodendrocytes at levels comparable to neurons^14^. If these molecules are sufficient to confer axonal synapses onto HEK293 cells, could a similar mechanism operate at the axon-oligodendrocyte interface? Specifically, could a similar mechanism operate to promote sheath growth and stability?

To identify genes that express synaptogenic proteins in oligodendrocytes, we queried both published transcriptome databases^14–16^ and in-house RNA-seq of fluorescence-activated cell sorted *mbp*+, *olig2*+ myelinating oligodendrocytes from *Tg(mbpa:tagRFPt; olig2:eGFP)* larvae^29^. We then selected six candidates based on oligodendrocyte expression (Fig. 4A,B) and the existence of published dominant negatives that disrupt synapse formation when expressed in neurons: Cadm1b (SynCAM1), Nlgn1 and Nlgn2b (Neuroligin1 and −2), Lrrc4ba (NGL-3), and Lrrtm1 and Lrrtm2 (Leucine rich repeat transmembrane proteins 1 and −2). We generated dominant-negative alleles of zebrafish proteins predicted to disrupt PDZ binding^27,30,31^. For all candidates, this included omitting conserved C-terminal PDZ binding motifs, encoding amino acids KEYYI in *cadm1b*^32^, ETQI in *lrrc4ba*^28^, STTRV in *nlgn1/2b*^30,33^, and ECEV in *lrrtm1/2*^27^. For *lrrtm1/2*, we omitted additional C-terminal residues (deleted last 55 AAs, including ECEV) to make alleles similar to those used by Linhoff et al (2009)^27^.

**Figure 4.**
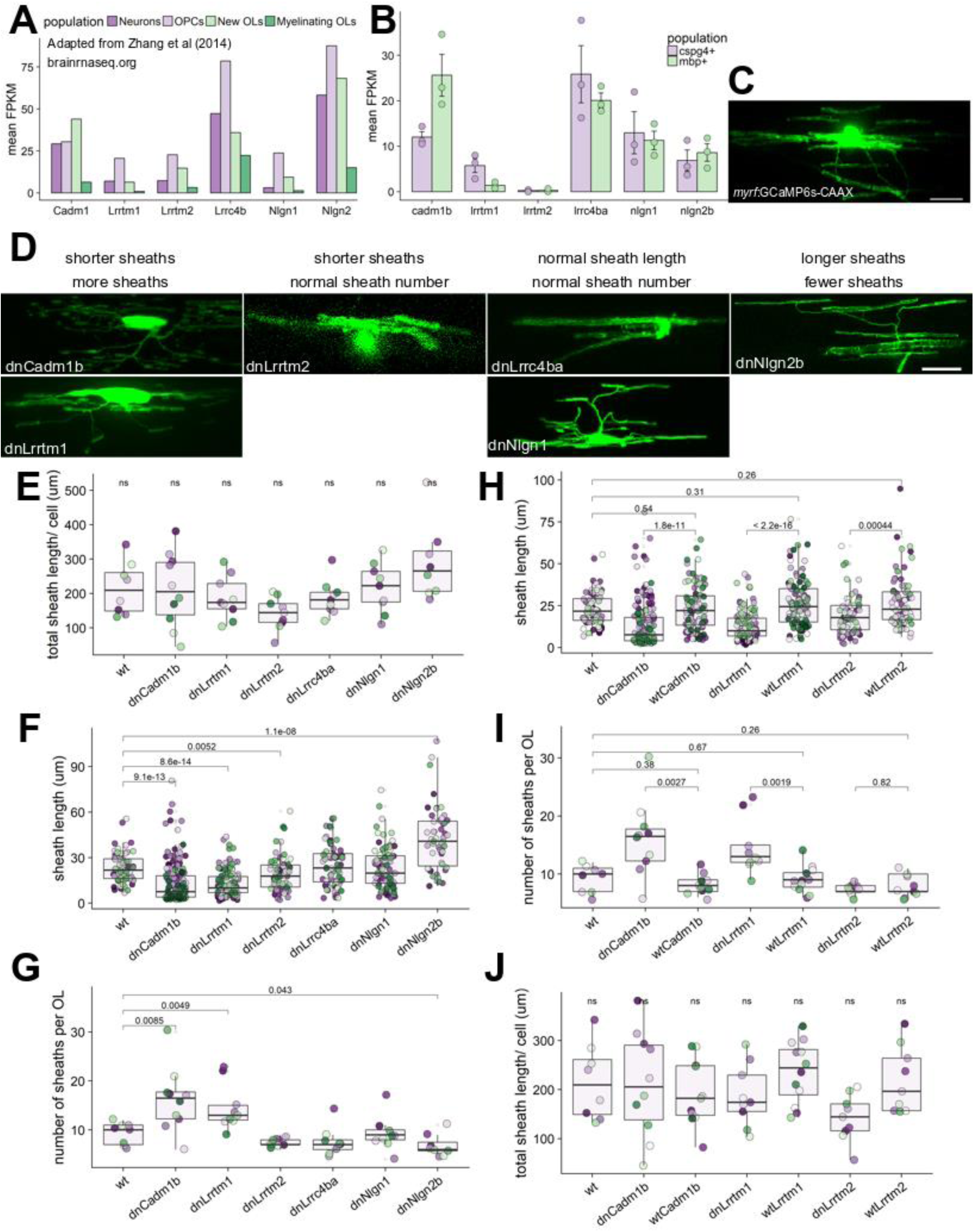
Candidate synaptogenic adhesion molecules have variable effects on myelin sheath length and number. (A) RNA-seq FPKM values for mouse cortical neurons, OPCs, new oligodendrocytes, and myelinating oligodendrocytes for candidates *Cadm1*, *Lrrtm1*, *Lrrtm2*, *Lrrc4b*, *Nlgn1*, and *Nlgn2*. Plot generated using publicly available values from brainrnaseq.org (Zhang et al, 2014). (B) RNA-seq FPKM values for FAC-sorted zebrafish *olig2*+/*cspg4*+ (cspg4+) and *mbpa*+/*olig2*+ (mbp+) cells for zebrafish homologs of the same candidates29. (C) Oligodendrocyte transiently expressing *myrf*:GCaMP6s-CAAX. Scale bar 10 µm. (D) Candidate dominant-negative (dn) alleles expressed transiently as *myrf*:dnX-2A-GCaMP6s-CAAX and grouped by effect on sheath length and number. Scale bar 10 µm. (E) Total sheath length per cell expressing each dominant-negative candidate was unchanged by all candidates, Kruskal-Wallis test. (F) Sheath length values for each dominant-negative candidate. n (cells/sheaths) = 8/74 wt, 10/159 dnCadm1b, 9/132 dnLrrtm1, 9/67 dnLrrtm2, 9/68 dnLrrc4ba, 9/83 dnNlgn1, 8/55 dnNlgn2b. Data in plots F-I were tested by Wilcox rank-sum with Bonferroni-Holm correction for multiple comparisons. dnLrrc4ba/wt and dnNlgn1/wt comparisons were ns. (G) Sheath number values for each dominant-negative candidate (n listed in (F)). (H) Decreased sheath length in dnCadm1b, dnLrrtm1, and dnLrrtm2 cells is specific to the DN disruption, because expression of the full-length (wt) protein does not reduce sheath length. n(cells/sheaths) = 11/91 wtCadm1b, 12/108 wtLrrtm1, 9/72 wtLrrtm2. I) Expression of WT forms of the three candidates that reduced sheath length resulted in normal sheath number. (J) Total sheath length per cell expressing each WT candidate was not significantly different for any group, Kruskal-Wallis test.

We expressed each of these dominant-negative constructs under the control of *myrf* regulatory DNA to disrupt transsynaptic adhesion specifically in oligodendrocytes. To label sheaths, we bicistronically expressed GCaMP6s-CAAX, which we made by fusing a CAAX motif to the C-terminus of GCaMP6s to promote membrane targeting. We chose to label cells with GCaMP6s-CAAX because it illuminates whole sheaths exceedingly well for sheath measurements. We imaged living larvae expressing each of the constructs, or a control *myrf*:GCaMP6s-CAAX (wt) construct (Fig. 4C,D), at 4 dpf to measure sheath number and length. The total sheath length generated per cell, a general measure of myelinating capacity, did not differ between any of the groups (Fig. 4E). However, oligodendrocytes expressing dnCadm1b, dnLrrtm1, and dnLrrtm2 formed significantly shorter sheaths, whereas oligodendrocytes expressing dnNlgn2b made longer sheaths (Fig. 4F). Intriguingly, dnCadm1b and dnLrrtm1 oligodendrocytes also elaborated several more sheaths than wildtype (wt), whereas dnNlgn2b oligodendrocytes generated slightly fewer sheaths (Fig. 4G). Because total sheath length per cell was unchanged, this suggests that these adhesion proteins do not influence the ability of oligodendrocytes to generate myelin but rather tune how oligodendrocytes allocate myelin among sheaths. Importantly, for all three dominant-negative constructs that reduced sheath length (dnCadm1b, dnLrrtm1, dnLrrtm2), expression of the full-length (wt) protein did not reduce sheath length relative to wt control (Fig. 4H), indicating that sheath-shortening is specific to disruption of PDZ binding for each of these proteins. Expression of full-length proteins also led to a normal number of sheaths generated per cell (Fig. 4I) and normal total membrane per cell (Fig. 4J). These data illustrate that a number of transsynaptic adhesion molecules, typically studied only in the context of neuron-neuron signaling at synapses, also determine the length and number of myelin sheaths formed by oligodendrocytes. Because conduction velocity along axons depends on myelin sheath length^10,34^, the reduction of myelin sheath length observed for dnCadm1b, dnLrrtm1, and dnLrrtm2 are likely to delay or otherwise disrupt the timely propagation of impulses within circuits.

### Cadm1b signals transsynaptically at the axon-myelin junction to promote ensheathment

Our dominant-negative approach tested the requirement for each candidate’s PDZ binding motif in modulating myelination. PDZ binding is essential for downstream signaling through postsynaptic scaffolds, including PSD-95 and CASK, in the recipient postsynaptic cell (or sheath). To determine where in the oligodendrocytes these candidates act to modulate ensheathment, we focused on one candidate, Cadm1b, due to the severity of the dominant-negative phenotype. We first confirmed expression of *cadm1b* in oligodendrocytes by generating a knock-in transgenic, using the same CRISPR/Cas9-mediated enhancer trap strategy for *cadm1b* as we did for *dlg4b* (Fig. 3A). By crossing *Tg(cadm1b:GAL4; UAS:eGFP-CAAX)* and *Tg(mbpa:tagRFP)* adults, we generated larvae expressing eGFP-CAAX in *cadm1b*+ cells and tagRFP in *mbp*+ cells (Fig. 5A). We found that most *mbp*+ cells are also *cadm1b*+ but not all *cadm1b*+ cells are *mbp*+, likely indicating neuronal expression.

**Figure 5.**
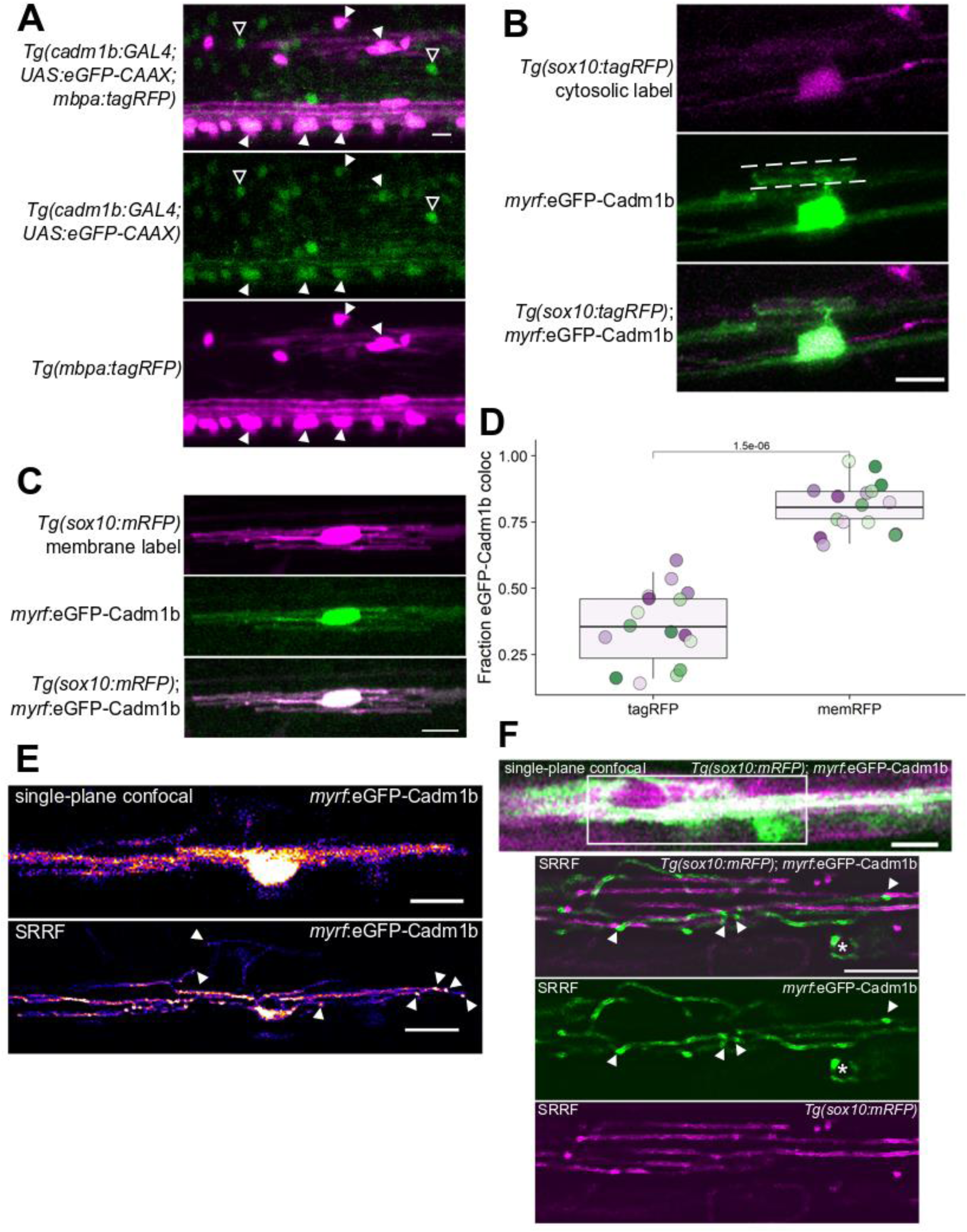
Cadm1b localizes to myelin sheath membrane. (A) CRISPR/Cas9-mediated GAL4 enhancer trap of *cadm1b* reports cells expressing *cadm1b* via *UAS*:eGFP-CAAX expression. Larvae additionally carrying *Tg(mbp:tagRFP)* reveal *mbp*+ myelinating oligodendrocytes. Closed arrowheads indicate *mbp*+, *cadm1b*+ oligodendrocytes, whereas open arrowheads indicate *mbp*-, *cadm1b*+ cells (likely neurons). Scale bar, 10 µm. (B, C) Expression of *myrf*:eGFP-Cadm1b in *Tg(sox10:tagRFP)* (B) or *Tg(sox10:mRFP)* (C) co-labeled oligodendrocytes. Dashed lines in (B) indicate parallel line-structure of eGFP-Cadm1b labeling. (D) Fraction of eGFP-Cadm1b colocalized with RFP (Mander’s colocalization coefficient) for *Tg(sox10:tagRFP)* (tagRFP) and *Tg(sox10:mRFP)* (memRFP). n=16 cells (tagRFP),16 cells (memRFP), Wilcox rank-sum test. (E) Conventional confocal single-plane image of an oligodendrocyte expressing eGFP-Cadm1b (top) and super resolution radial fluctuations (SRRF) processing of 197 single-plane frames of the same cell (bottom). Arrowheads indicate puncta of eGFP-Cadm1b signal. Scale bars, 10 µm. (F) Confocal single-plane (top) and SRRF (bottom 3 panels) imaging of a eGFP-Cadm1b expressing oligodendrocyte in a *Tg(sox10:mRFP)* larva. Arrowheads indicate eGFP-Cadm1b puncta that are not present in mRFP SRRF. Asterisk marks a sheath going in to the plane of view with circular edges surrounded by eGFP-Cadm1b puncta. Scale bars, 5 µm.

To track Cadm1b localization in oligodendrocytes, we generated a fusion protein, eGFP-Cadm1b, and drove expression with *myrf* regulatory DNA. We found eGFP-Cadm1b distributed in both the somatic and sheath compartments (Fig. 5B). In sheaths, eGFP-Cadm1b signal resembled two parallel lines demarcating the sheath perimeter, raising the possibility that eGFP-Cadm1b is transmembrane in sheaths. To estimate the fraction of our fusion protein associated with sheath membrane vs cytosol, we expressed eGFP-Cadm1b in oligodendrocytes that were transgenically co-labeled with either *Tg(sox10:tagRFP)*, which expresses a cytosolic RFP (Fig. 5B), or *Tg(sox10:mRFP)*, which expresses a membrane-tethered RFP (Fig. 5C). We calculated Mander’s colocalization coefficient, the fraction of eGFP+ points that are also RFP+, using the Fiji plugin JACoP (Just Another Colocalization Plugin)^35^. We found that 35.3% ± 3.4% of eGFP-Cadm1b signal was colocalized with cytosolic tagRFP, whereas 81.1% ± 2.2% of eGFP-Cadm1b signal was colocalized with membrane RFP (Fig. 5D), indicating that eGFP-Cadm1b is membrane-localized in sheaths.

Does Cadm1b exhibit sub-sheath localization, similar to the axonal placement of SypHy hotspots (Fig. 2C) and PSD-95 puncta at the terminal ends of sheaths (Fig. 3D)? To obtain a more detailed view of eGFP-Cadm1b localization in sheaths, we used an analytical method for super-resolution imaging, super resolution radial fluctuations (SRRF)^36^. SRRF revealed eGFP-Cadm1b puncta primarily at the terminal ends of sheaths (Fig. 5E, bottom). To determine whether SRRF would report this labeling pattern for any membrane-associated protein, we performed SRRF imaging of cells expressing both *Tg(sox10:mRFP)* and *myrf*:eGFP-Cadm1b (Fig. 5F). We found that most eGFP-Cadm1b puncta are not associated with mRFP puncta, suggesting that eGFP-Cadm1b enrichment at sheath ends is a specific feature of Cadm1b rather than a general feature of membrane-associated proteins. Our observations that most axonal exocytosis sites are punctate and located near terminal ends of sheaths coupled with the frequent terminal localization of PSD-95 puncta raises the possibility that the synaptic features of the axon-myelin interface are primarily present at the terminal ends of sheaths.

This sheath membrane localization is consistent with Cadm1b functioning in sheaths as a transmembrane protein, possibly bridging the axon-myelin “cleft” to interact with the axon. To test whether Cadm1b specifically interacts with an extracellular partner located on the axon, we generated a second dominant-negative allele designed to prevent adhesion to transsynaptic partners. This allele, Ig1-dnCadm1b, lacks the extracellular distalmost Ig-like domain (Ig1) that specifically interacts transsynaptically with Cadm partners located on other cells^37^. We excised Ig1 while preserving the rest of the extracellular domain because the proximal Ig-like domains (Ig2, Ig3) allow Cadm1b to be incorporated into *cis* oligomers with other Cadm1b molecules. In this way, Ig1-dnCadm1b is predicted to bind endogenous Cadm1b via lateral Ig2 and Ig3 interactions, but to interfere with adhesion in *trans*^37^.

Expression of Ig1-dnCadm1b in oligodendrocytes produced a phenotype distinct from expression of the wt form or the PDZ binding dominant-negative (hereafter called PDZIIb-dnCadm1b) (Fig. 6A-A”). Ig1-dnCadm1b oligodendrocytes generated sheaths that were significantly shorter than wt and wtCadm1b-expressing oligodendrocytes but were longer than PDZIIb-dnCadm1b oligodendrocytes (Fig. 6B). Furthermore,the number of sheaths generated was modestly increased, but not significantly different that wt or wtCadm1b sheath number (Fig. 6C). The myelinating capacity of cells expressing each of the constructs was unchanged (Fig. 6D), indicating that both extracellular adhesion and PDZ binding tune how oligodendrocytes distribute myelin among sheaths rather than influencing myelin production. Because PDZIIb-dnCadm1b specifically disrupts downstream Cadm1b interactions with scaffolds including CASK, syntenin, and Mpp3^26,38^, whereas Ig1-dnCadm1b disrupts extracellular adhesion without changing PDZ interactions^37^, these data suggest that Cadm1b has roles in both “postsynaptic” signaling within sheaths as well as transsynaptic adhesion-induced signaling with the axon to tune ensheathment, perhaps via induction of presynaptic assembly in the axon (Fig. 7).

**Figure 6.**
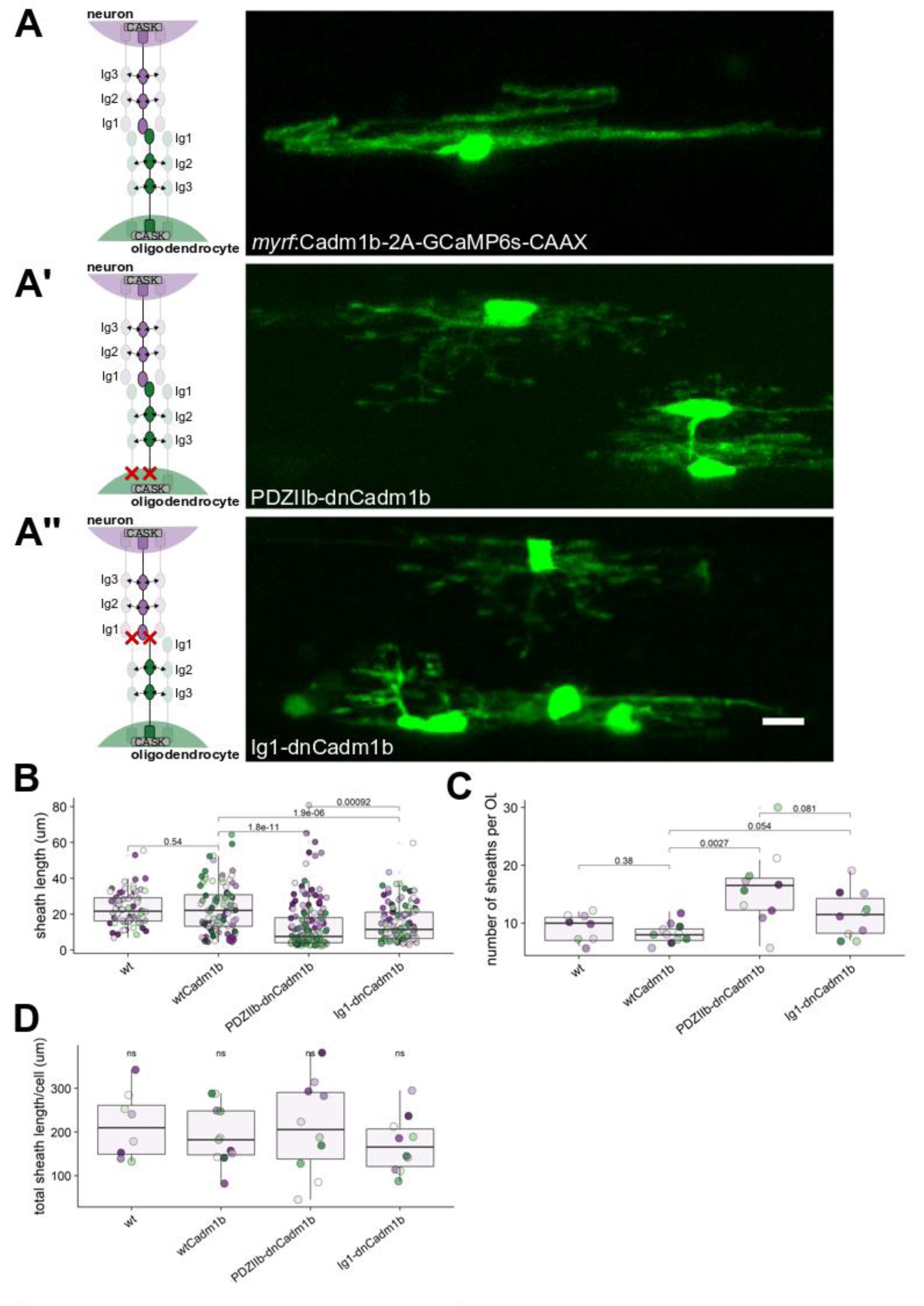
The extracellular, *trans*-acting adhesion domain (Ig1) of Cadm1b promotes myelin sheath growth. (A) Left, schematic of oligodendrocyte Cadm1b (green) interacting with partner Cadm1/2/3/4 located on neurons. Right, oligodendrocyte overexpressing wildtype Cadm1b with bicistronic GCaMP6s-CAAX to label sheath membrane. Scale bar, 10 µm. (A’) Left, schematic of oligodendrocyte PDZIIb-dnCadm1b, which is predicted to interfere with binding to PDZ scaffolds including CASK, Syntenin, and Mpp3. Right, oligodendrocytes expressing PDZIIb-dnCadm1b. Scale bar, 10 µm. (A’’) Similar to (A), (A’) for allele Ig1-dnCadm1b, which lacks the extracellular Ig1 domain and is predicted to disrupt extracellular adhesion. (B) Sheath lengths for wt, wtCadm1b-, PDZIIb-dnCadm1b-, and Ig1-dnCadm1b-expressing oligodendrocytes. Note that PDZIIb-dnCadm1b is the same allele presented in Figure 4. n (cells/sheaths) = 8/74 (wt), 11/91 (wtCadm1b), 10/159 (PDZIIb-dnCadm1b), 10/115 (Ig1-dnCadm1b), Wilcox rank-sum test with Bonferroni-Holm correction for multiple comparisons. (C) Sheath number for wt, wtCadm1b-, PDZIIb-dnCadm1b-, and Ig1-dnCadm1b-expressing oligodendrocytes. Same n and statistical test as in (B). (D) Total sheath length generated per cell is unchanged by all alleles, Kruskal-Wallis test.

**Figure 7.**
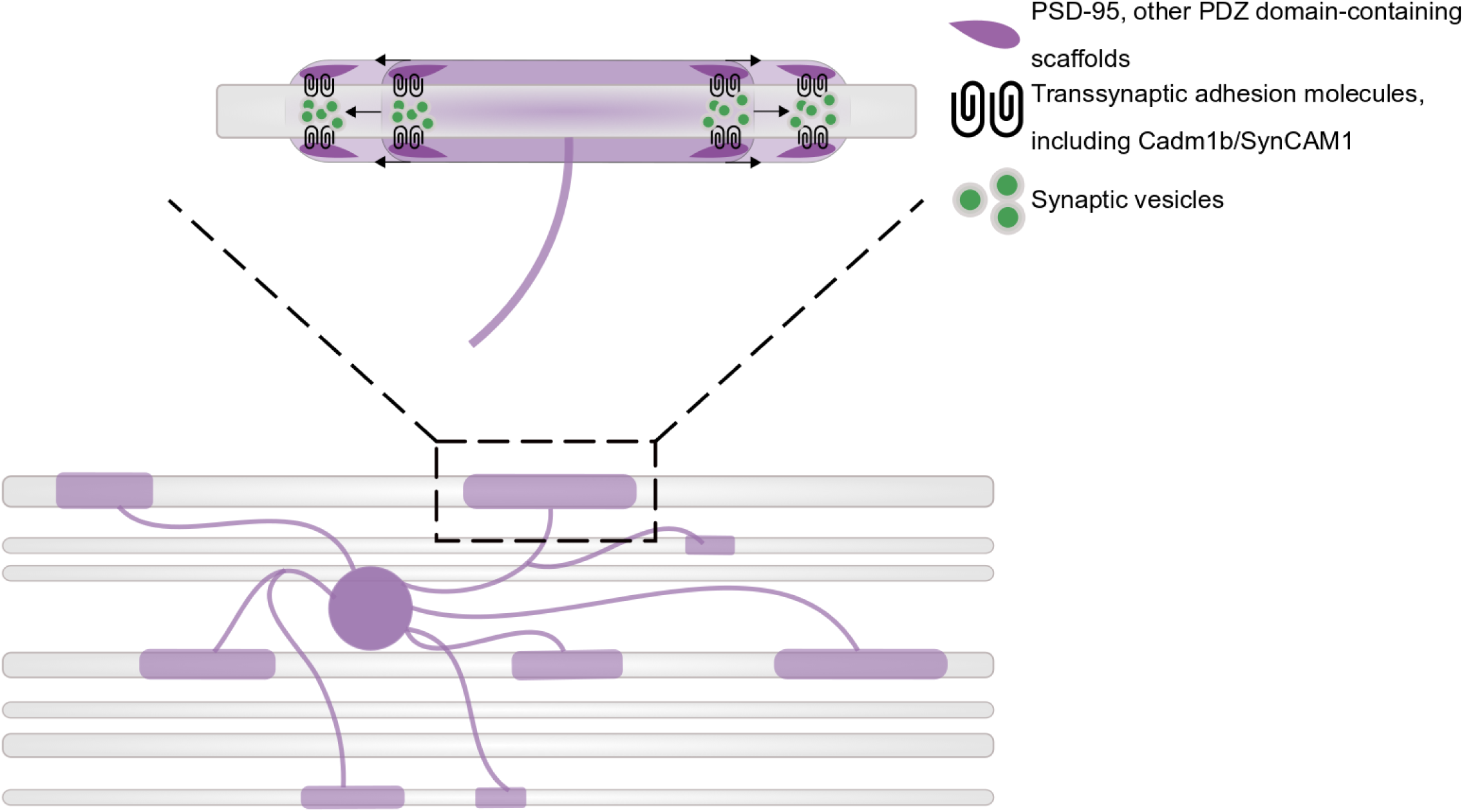
Working model of adhesion-promoted sheath growth. PSD-95 and other PDZ domain-containing scaffolds expressed by oligodendrocytes, including CASK and Dlg1, anchor transsynaptic adhesion molecules that adhere to and induce presynaptic assembly in the ensheathed axon. The accumulation and exocytosis of synaptic vesicles at ensheathment sites promotes the elongation of nascent sheaths, possibly via neurotransmitter- and/or neurotrophin-induced intracellular signaling in the sheath.

## DISCUSSION

Our work supports a model of activity-regulated myelination that is mechanistically similar to synapse formation. We investigated this similarity because both dendrites and myelin sheaths share an early dependence on synaptic vesicle release as well as expression of many key regulators of postsynaptic maturation. We found components of vesicular release machinery, Synaptophysin and Synaptobrevin, located under myelin sheaths that form on *phox2b*+ axons. This machinery is functional, as alkaline trapping of the pH-sensitive reporter SypHy reported sites of exocytosis located under myelin sheaths. On the “postsynaptic” side of this interaction, we found localization of the postsynaptic scaffolding molecule PSD-95 in sheaths by using a genetically-encoded intrabody directed to endogenous PSD-95.

Oligodendrocytes previously have been shown to express the postsynaptic scaffold PSD-95 by immunoblot^39^ and RNA-seq^14–16^, but understanding how oligodendrocytes use this scaffold required a method that can resolve the subcellular localization of the protein. We used a genetically-encoded intrabody to label endogenous PSD-95 in oligodendrocytes and discovered heterogeneity in the labeling of sheaths of individual oligodendrocytes. Some sheaths exhibited end labeling only, while others exhibited a periodic spread of puncta along the length of the sheath. Still other sheaths exhibited no defined labeling. Why this diversity? Perhaps sheaths, like dendritic spines, use different scaffolds to anchor unique subsets of receptors and signaling molecules required for autonomous interaction with the presynaptic axon. For example, excitatory synapses use PSD-95 to anchor NMDA receptors, whereas inhibitory synapses primarily use Gephryin (Gphn) to anchor GABA receptors^40^. Oligodendrocytes also express Gphn and other scaffolds^14^, raising the possibility that sheaths labeled poorly for PSD-95 might instead utilize other protein scaffolds, perhaps to support sheaths on different classes of neurons.

Why have presynaptic and postsynaptic machinery not been discovered at the axon-myelin interface by transmission electron microscopy (EM), a technique that has been used to identify synapses for over 30 years? One possibility for this lapse is that EM identifies mature synapses, but frequently misses small, nascent, and diverse synapses that do not resemble classic EM synapses^41^. Wake et al (2015) previously used EM to assess axon-OPC contacts *in vitro*. While they observed presynaptic vesicles docked at axon-OPC contacts, they did not identify a postsynaptic density, which led them to conclude that the junction is nonsynaptic^13^. In contrast, by using a non-EM approach, we have detected PSD-95 in sheaths. What explains this difference? Perhaps axon-OPC junctions have PSDs on par with immature synapses, which lack an EM-resolvable PSD until maturity, or perhaps maturity of axon-OPC junctions was not achieved in culture. Another possible reason that synaptic features of axon-oligodendrocyte contacts may go unnoticed by EM is diversity: perhaps synaptic features are not equally present at all axon-oligodendrocyte junctions. Like Wake et al (2015), Doyle et al (2017) identified docked axonal vesicles under myelin, but not all myelinated axons had sub-myelin docked vesicles^42^. This is consistent with the observation that vesicular release from certain axons profoundly modulates their myelin profiles, while vesicular release is dispensable for the myelination of other classes of axons^43^. Furthermore, while we found PSD-95 localization in most sheaths, there was substantial heterogeneity between sheaths of individual cells. Together, these studies raise the possibility that axon-oligodendrocyte contacts are diverse in their usage of synaptic elements, perhaps as broad in scope as the variety of synapses. This prompts a significant need for alternatives to EM to identify diverse synapses. Burette et al (2015) set out recommendations for single-synapse analysis for those synapses missed by EM, including fluorescence microscopy and optophysiology approaches^41^, both of which we have used here.

We sought to test whether these synaptic similarities are integral to normal myelination or merely inconsequential features of this junction. Because activity manipulations fail to prevent CNS synapse formation^44–46^, and our lab’s previous work indicates that tetanus toxin does not prevent ensheathment^11^, we instead interrupted an upstream director of synapse formation, cell adhesion, to test whether manipulating synaptic adhesion systems would negatively impact myelination via interruption of synaptic features of the interface. We screened dominant-negative versions of candidate transsynaptic adhesion molecules expressed by oligodendrocytes and measured oligodendrocyte sheath length, number, and total sheath length per cell. In doing so, we discovered that oligodendrocytes may use multiple adhesion molecules in tandem to tune sheath length and number. At synapses, multiple transsynaptic adhesion systems are thought to operate in parallel^47^, and redundancy within and between families can obscure individual contributions. Despite possible redundancy, we unveiled specific requirements for PDZ binding of Cadm1b, Lrrtm1, and Lrrtm2 in oligodendrocyte sheath length. Lrrtm1 and Lrrtm2 bind to the PDZ domains of PSD-95^27^, but Cadm1b is instead anchored by the related scaffold CASK, which also anchors the major oligodendrocyte protein Claudin11^48^.

We pursued Cadm1b due to the severity of oligodendrocyte morphology in our screen: disruption of PDZ-binding reduced sheath length by nearly 50%. Shorter sheaths are predicted to reduce the conduction velocity of ensheathed axons^34^. However, oligodendrocytes also serve physiological functions uncoupled from morphology. An oligodendrocyte-specific Kir4.1 knockout mouse has overtly normal oligodendrocytes and myelin, but animals exhibit ataxia and spontaneous seizures due to poor K+ uptake by oligodendrocytes^49^. An important lesson from this work is that oligodendrocyte manipulations can have drastic effects on CNS health without presenting morphological defects. Two candidates in our screen, dnLrrc4ba and dnNlgn1, were not significantly different from wt oligodendrocytes in any morphological parameter we assessed. However, we cannot conclude that these transsynaptic adhesion molecules are dispensable in oligodendrocytes. Neurexin-neuroligin adhesion can induce axon-oligodendrocyte adhesion *in vitro*^50^, and Lrrc4 may modulate cell cycle progression in glia^51^. Furthermore, redundancy and compensation within adhesion protein families may obscure the function of individual adhesion proteins. For example, NLGN1, −2, −3 triple knockout mice exhibit functional synaptic deficits and perinatal lethality that are not present in single and double knockout animals^52^. Oligodendrocyte-specific deletion or mutation of the genes encoding these candidates in a model permitting physiological and behavioral assays may be required to determine the function of these proteins in myelination.

We have shown that synaptic features are present at the axon-myelin junction and that synaptogenic adhesion molecules have a previously underappreciated role in shaping myelination. Is this similarity to synaptogenesis biologically significant? Dendrites and oligodendrocytes utilizing similar mechanisms to establish stable contacts with axons raises the possibility that oligodendrocytes may be vulnerable to pathology in psychiatric disease, where mutations in synaptic genes are cited as likely drivers of pathogenesis^53^. Indeed, ultrastructural analysis of postmortem brain tissue from patients with schizophrenia reveals myelin and oligodendrocyte defects^54^ and diffusion tensor imaging of patients living with schizophrenia^55^ and autism^56^ reveal white matter defects. Together, these findings raise the intriguing possibility that mutations in synaptic genes could affect oligodendrocytes cell-autonomously to disrupt myelination, which may then contribute to disease progression via reduced neuronal support or conduction velocity. Alternatively, mutations in synaptic genes could impair synaptic transmission and thus change neuronal activity in a way that disrupts oligodendrocyte maturation, myelination, or survival. We have shown that a number of synaptic proteins are expressed at the axon-myelin interface and that disruption of some of these proteins in oligodendrocytes disrupts myelination. This finding supports the former possibility, that synaptic genes cell-autonomously regulate myelination in oligodendrocytes. However, because the oligodendrocyte lineage is sensitive to perturbations in neuronal activity^9^, it is also possible that reduced myelination resulting from synaptic protein dysfunction in oligodendrocytes causes aberrant or asynchronous neuronal activity which then further disrupts the oligodendrocyte lineage. Untangling the relationship between neuronal activity and oligodendrocyte myelination is crucial for understanding nervous system function in both health and disease. Our work draws structural and functional parallels between synaptogenesis and myelination and provides new targets for investigating the mechanistic basis of activity-regulated myelination.

## Methods

### Zebrafish lines and husbandry

All animal work was approved by the Institutional Animal Care and Use Committee at the University of Colorado School of Medicine. Zebrafish embryos were raised at 28.5°C in embryo medium and staged according to hours or days post fertilization (hpf/ dpf) and morphological criteria^57^. *Tg(sox10:mRFP)*^*vu234*^, *Tg(sox10:tagRFP)*^*co26*^, *Tg(phox2bb:GAL4)*^*co21*^, *Tg(UAS:syp-eGFP)* ^58^*, Tg(mbpa:tagRFP)*^*co25*^, and *Tg(UAS:eGFP-CAAX)*^*co18*^ transgenic lines were used. Additionally, we generated and used the new lines *Tg(cadm1b:GAL4)*^*co53*^ and *Tg(dlg4b:GAL4)*^*co54*^ (see section on enhancer trapping). All other reporters were expressed by transient transgenesis to achieve sparse labeling for single cell analysis.

### Plasmid construction and generation of transgenic zebrafish

Tol2 expression plasmids were generated by Multisite Gateway cloning and injected into 1-cell embryos with Tol2 mRNA to generate transient transgenic animals. The following entry clones were used (cited) or made (see table of primers) via BP recombination of appropriate pDONR backbone with attB-flanked regulatory elements or coding sequences.

*p5E-neuroD*^59^*, p5E-4xUAS*^60^*, p5E-myrf* (gift from Jacob Hines)

*pME-eGFP*^60^*, pME-GAL4*^60^*, pME-vamp2, pME-sypHy, pME-GCaMP6s-CAAX, pME-cadm1b, pME-Ig1-dnCadm1b, pME-PDZIIb-dnCadm1b, pME-lrrtm1, pME-dnLrrtm1, pME-lrrtm2, pME-dnLrrtm2, pME-dnLrrc4ba, pME-dnNlgn1, pME-dnNlgn2b, pME-PSD95.FingR-GFP-CCR5TC-KRAB(A)*

*p3E-polyA*^60^*, p3E-eGFP*^60^*, p3E-2A-GCaMP6s-CAAX, p3E-cadm1b*

Entry clones were LR-recombined with either of destination vectors *pDEST-Tol2-CG2* (green heart marker) or *pDEST-Tol2-CC2* (blue eye marker) as transgenesis indicators^60^.

*zcUAS:PSD95.FingR-GFP-CCR5TC-KRAB(A)* was a gift from Joshua Bonkowsky (Addgene plasmid # 72638) and was used to generate *pME-PSD95.FingR-GFP-CCR5TC-KRAB(A)*. *CMV::SypHy A4* was a gift from Leon Lagnado (Addgene plasmid # 24478) and used to generate *pME-sypHy*.*pGP-CMV-GCaMP6s* was a gift from Douglas Kim (Addgene plasmid # 40753) and used to generate *pME-GCaMP6s-CAAX* and *p3E-2A-GCaMP6s-CAAX*.

### CRISPR/Cas9-mediated enhancer trapping

We knocked in *GAL4* 200-500 bp upstream of the translation start site of *cadm1b* and *dlg4b* using a previously published method^21^. Briefly, *mbait-hs-Gal4* plasmid, mBait guide RNA, guide RNA targeting upstream of our genes of interest (see primer table for sequences), and Cas9 mRNA were injected into 1-cell *Tg(UAS:eGFP-CAAX)* embryos. F0 founders were screened for expression and raised to adulthood. Experiments involving these knock-ins were only performed on stable transgenic (F1 or later) *Tg(cadm1b:GAL4)*^*co53*^ and *Tg(dlg4b:GAL4)*^*co54*^ larvae.

### Dominant negative allele generation

We made zebrafish homologs of published mouse dominant-negative alleles for all candidates. For candidates with duplicate paralogs in the zebrafish genome, we investigated the paralog with higher expression in oligodendrocytes^29^. We isolated RNA from whole, wildtype AB-strain 4 dpf larvae and synthesized cDNA (iScript) to use as template to high-fidelity (Phusion) PCR amplify attB-flanked coding sequences for candidates, omitting C-terminal PDZ binding sites for Cadm1b (-KEYYI)^31,32^, Lrrtm1 and Lrrtm2 (-ECEV)^27^, Nlgn1 and Nlgn2b (-STTRV)^30^, and Lrrc4ba (-ETQI)^27^. Full-length sequences were also amplified. AttB-flanked products were recombined with *pDONR-221* to form middle entry vectors, and subsequently recombined with *p5E-myrf*, *p3E-2A-GCaMP6s-CAAX*, and *pDEST-Tol2-CG2* to generate expression constructs.

### Statistics

All statistics were performed in R (version 3.4.1) with RStudio. Plots were generated using dplyr and ggplot2 packages^61^ with cowplot package formatting^62^, and all statistical tests were performed using ggpubr^63^ except for the Chi square test, which was carried out in base R^64^. We used the Wilcox rank sum test (also called Mann-Whitney), with no assumption of normality, for all unpaired comparisons. For multiple comparisons, we first assessed global significance using the Kruskal-Wallis test, followed (only if Kruskal-Wallis significant) by pairwise Wilcox rank sum tests with Bonferroni-Holm correction for multiple comparisons.

### Image analysis

We performed all imaging on live larvae, embedded laterally in 1.2% low-melt agarose containing either 0.4% tricaine or 0.3 mg/ml pancuronium bromide for immobilization. We acquired images on a Zeiss Axiovert 200 microscope with a PerkinElmer spinning disk confocal system (PerkinElmer Improvision) or a Zeiss LSM 880 (Carl Zeiss). After collecting images with Volocity (PerkinElmer) or Zen (Carl Zeiss), we performed all processing and analysis using Fiji/ImageJ.

### Synaptophysin-eGFP clustering

We imaged *Tg(phox2b:GAL4; UAS:syp-eGFP; sox10:mRFP)* larvae at 3, 4, and 5 dpf. Roughly 50% of *phox2b+* neurons are myelinated^11^, so we confirmed myelination status of individual *phox2b*+ neurons by taking oversampled z-stacks and examining orthogonal views. We traced individual *phox2b*+ neurons containing Syp-eGFP puncta with Fiji’s Plot Profile tool and counted peaks, representing individual puncta, that exceeded 150% of the inter-peak background fluorescence intensity. The number of peaks (puncta) per µm of ensheathed and bare axon were transferred from Fiji to R and density of puncta in both regions was determined by dividing puncta count by length (for bare axon density, we used the length of axon present in the field of view). Regions were compared age-wise by Wilcox rank-sum test (wilcox.test, R package ggpubr) with no assumption of normality.

### SypHy signatures under myelin sheaths

*neuroD*:sypHy; *Tg(sox10:mRFP)* larvae were immersed for 1 hour in 1 µM bafilomycin A1 (Tocris, cat 1334) in DMSO vehicle in embryo medium. We then paralyzed embryos by adding pancuronium bromide (Sigma, cat P1918) to the solution to achieve a final concentration of 0.3 mg/ml. We made a small incision to the tip of the tail with a tungsten needle to promote circulation of the paralytic agent. Larvae were monitored for an additional 24h post-experiment to ensure incisions did not cause lasting injury or fatality.

We imaged larvae in pancuronium bromide-containing low-melt agarose immersed in the same treated embryo medium (containing bafilomycin and pancuronium). We acquired oversampled z-stacks at a single x-y position, over the yolk extension, for individual larvae (1 z-stack per fish). Determination of whether an axonal sypHy hotspot was wrapped by mRFP+ myelin was made by orthogonal views. Because category designation (punctate, filled, impartial) was subjective we plotted individual observations for transparency. Among the punctate-ensheathing category, the number of puncta was measured the same way Syp-eGFP density was assessed. Pearson’s correlation was calculated for the number of sypHy hotspots under each sheath vs sheath length using the stat_cor() function in the ggpubr R package.

### PSD-95 localization in sheaths

We imaged transient-transgenic *myrf*:PSD95.FingR-GFP-CCR5TC-KRAB(A); *Tg(sox10:mRFP)* and transient-transgenic *myrf*:GAL4; *zcUAS*:PSD95.FingR-GFP-CCR5TC-KRAB(A); *Tg(sox10:mRFP)* larvae for examination of unregulated and regulated PSD95.FingR expression in oligodendrocytes, respectively. Because category designation (end, periodic, diffuse) was subjective, observations for individual cells were plotted for transparency. Unique cells were examined at each time point; i.e., cell #1 at 3 dpf is not cell #1 at 4 dpf. Because sheaths overlapping in x-y space with the soma were frequently blown out, only those that did not overlap in x-y space with the soma were analyzed to prevent incorrect category assignment due to brightness. To test whether the distribution of categories changes over developmental time, we carried out a Chi square test (chisq.test, base R) on the contingency table for category at each age point.

### Dominant-negative tests and sheath measurements

All imaging was performed at 4 dpf at approximately the same time each day (∼100-102 hpf). We predetermined that a sample size of 9-12 cells per candidate, with no more than 2 cells per animal, would generate a sufficient number of sheaths for both sheath length and number measurements. Sheaths were measured per cell by linear ROIs captured in ImageJ’s ROI manager. Length values were exported to R with cell identification numbers to identify the number of lengths (sheaths) per cell. Total sheath length per cell is the sum of sheath lengths per cell identification number and sheath number is the number of lengths associated with an identification number. For all three parameters (total sheath length per cell, sheath length, and sheath number) we first tested for global significance using the Kruskal-Wallis test, and if this was significant we made pairwise comparisons between groups using the Wilcox rank-sum test with p-values adjusted for multiple comparisons by the Bonferroni-Holm method.

### Quantitative colocalization

We used the Fiji plugin JACoP (Just Another Colocalization Plugin)^35^ to calculate Mander’s Colocalization Coefficients, M1 and M2, which describe the fraction of protein A that colocalizes with protein B and vice versa, respectively^65^. We chose to evaluate colocalization on the basis of these coefficients because they are independent of fluorescence intensity, which varies between transient-transgenic cells and animals. Briefly, single plane images of oligodendrocytes expressing *myrf*:eGFP-Cadm1b on either a membrane-labeling *Tg(sox10:mRFP)* or cytosol-labeling *Tg(sox10:tagRFP)* background were cropped to minimize background and split into separate channels for thresholding. JACoP generated M1 and M2 values, which were then exported to R for analysis.

### Super resolution radial fluctuations (SRRF)

For individual cells, 100-200 single z-plane images were acquired with minimal time delay between images (acquisition rate ∼5 frames per second). Images were transferred from Zen to Fiji and first corrected for drift (Plugins > Registration > Correct 3D drift). We corrected for slow drifts but did not edge-enhance images to avoid introducing artefacts. Drift-corrected images were then analyzed using the NanoJ-SRRF plugin with default settings.

#### Table of primers/ other DNA oligonucleotides

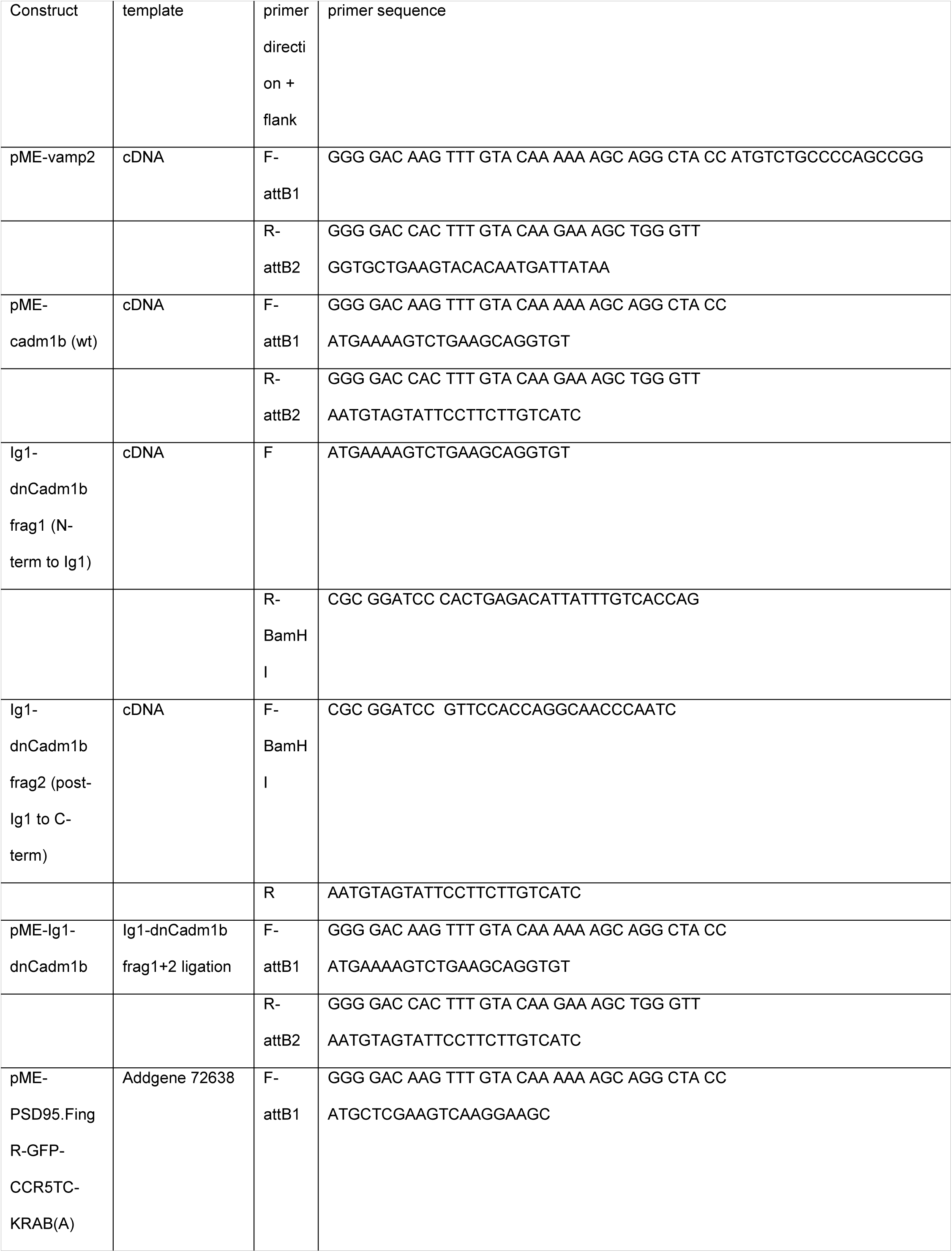

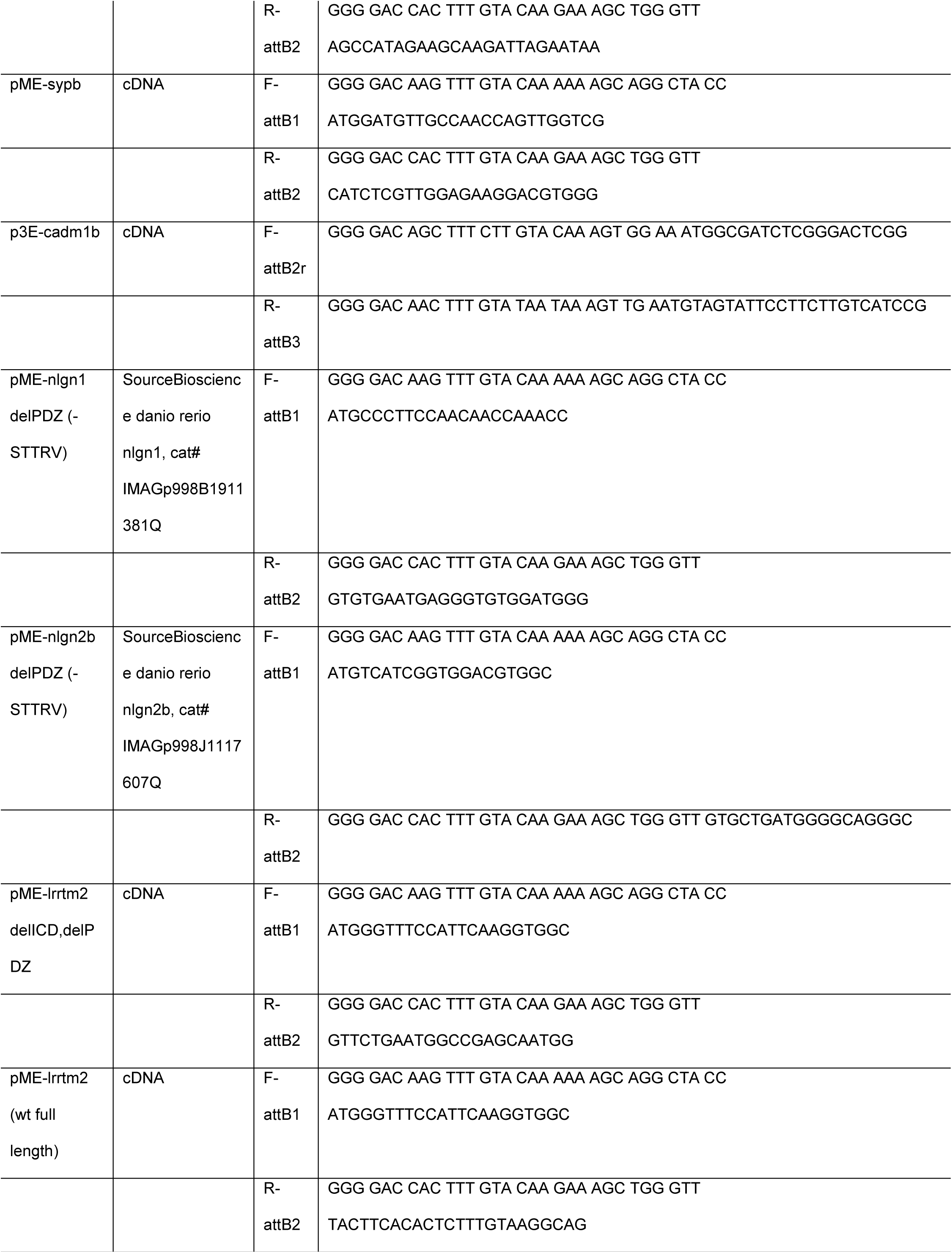

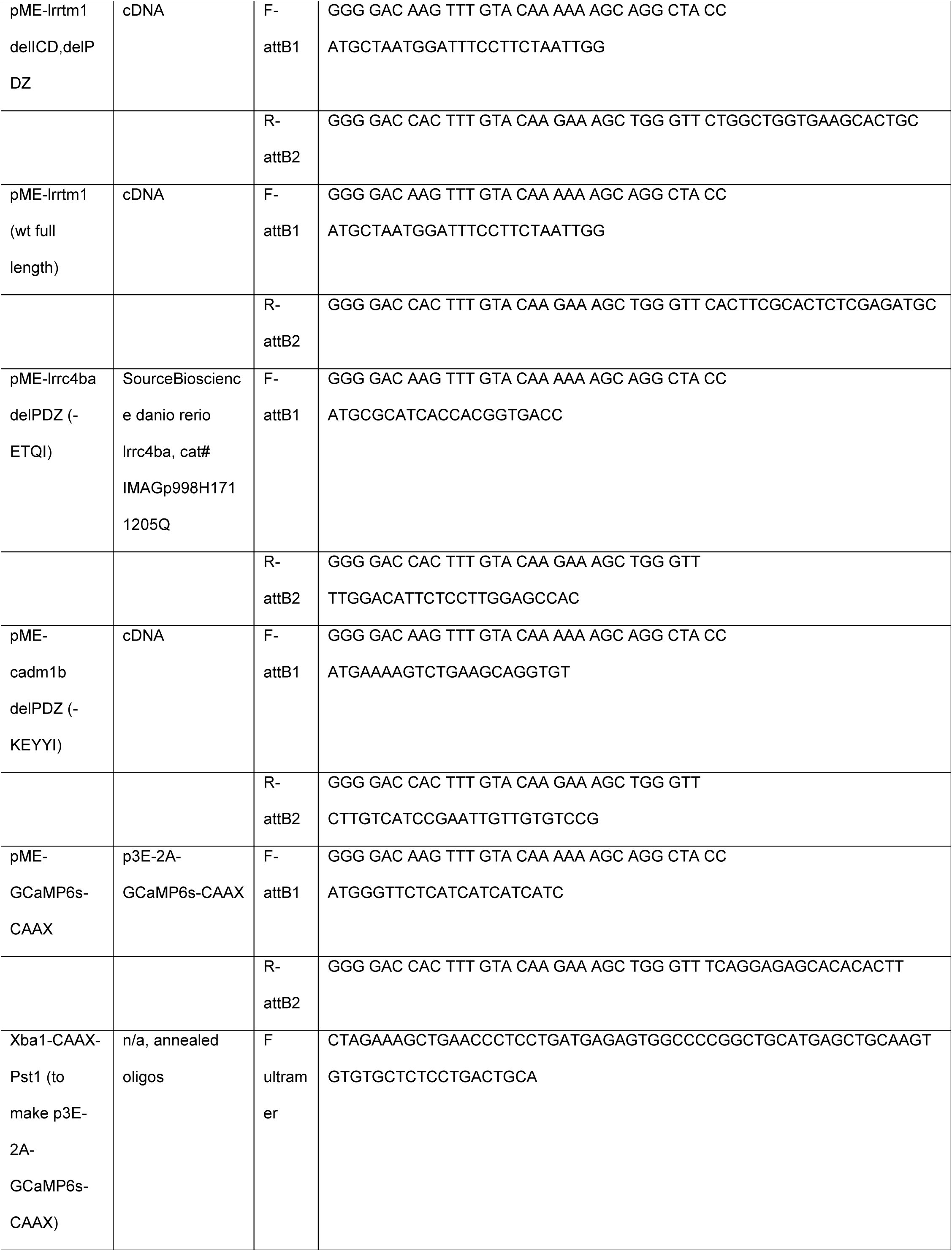

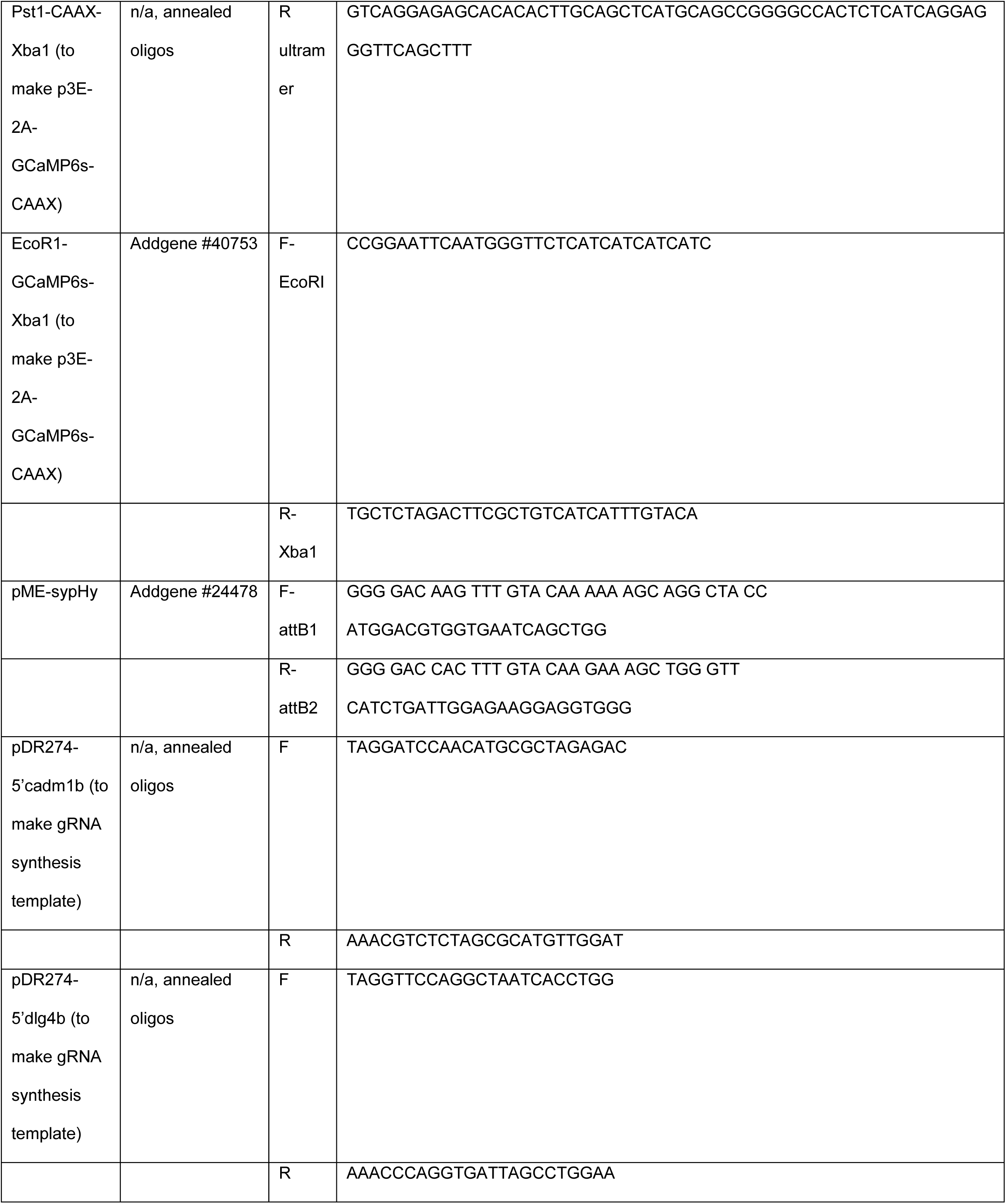

## ACKNOWLEDGMENTS

We are grateful to Jacob Hines for cloning *myrf* regulatory DNA and providing it to us. We also thank Ethan Hughes and Caleb Doll for valuable comments on the manuscript. This work was supported by US National Institutes of Health (NIH) grant R01 NS046668 and a gift from the Gates Frontiers Fund to B.A. and a National Science Foundation Graduate Research Fellowship (DGE-1553798) to A.N.H. The University of Colorado Anschutz Medical Campus Zebrafish Core Facility was supported by NIH grant P30 NS048154. All DNA plasmids and transgenic zebrafish used in this study are available by request.

## AUTHOR CONTRIBUTIONS

A.N.H. and B.A. conceived the project. A.N.H. performed all the experiments and collected and analyzed all the data. A.N.H. wrote and B.A. edited the manuscript.

